# Attenuation of homeostatic signaling from apoptotic thymocytes triggers a global regenerative response in the thymus

**DOI:** 10.1101/2020.08.31.275834

**Authors:** Sinéad Kinsella, Cindy A. Evandy, Kirsten Cooper, Lorenzo Iovino, Paul C. deRoos, Kayla S. Hopwo, David W. Granadier, Colton W. Smith, Shahin Rafii, Jarrod A. Dudakov

## Abstract

The molecular triggers of organotypic tissue repair are unknown. The thymus, which is the primary site of T cell development, is a model of tissue damage and regeneration as it is particularly sensitive to insult, but also has a remarkable capacity for repair. However, acute and profound damage, such as that caused by common cytoreductive therapies or age-related decline, lead to involution of the thymus and prolonged T cell deficiency, precipitating life-threatening infections and malignant relapse. Consequently, there is an unmet need to boost thymic function and enhance T cell immunity. Here, we demonstrate an innate trigger of the reparative response in the thymus, centered on the attenuation of signaling directly downstream of apoptotic cell detection as thymocytes are depleted after acute damage. We found that the intracellular pattern recognition receptor NOD2, via induction of microRNA-29c, suppressed the induction of the regenerative factors IL-23 and BMP4, from thymic dendritic cells (DCs) and endothelial cells (ECs), respectively. During steady-state, when a high proportion of thymocytes are undergoing apoptosis (as a consequence of selection events during T cell development), this suppressive pathway is constitutively activated by the detection of exposed phosphatidylserine on apoptotic thymocytes by cell surface TAM receptors on DCs and ECs, with subsequent downstream activation of the Rho GTPase Rac1. However, after damage, when profound cell depletion occurs across the thymus, the TAM-Rac1-NOD2-miR29c pathway is abrogated, therefore triggering the increase in IL-23 and BMP4 levels. Importantly, this pathway could be modulated pharmacologically by inhibiting Rac1 GTPase activation with the small molecule inhibitor EHT1864, leading to increased thymic function and T cell recovery after acute damage. In conclusion, our work not only represents a novel regenerative strategy for restoring immune competence in patients whose thymic function has been compromised due to cytoreductive conditioning, infection, or age; but also, identifies a mechanism by which tissue regenerative responses are triggered.

## INTRODUCTION

Efficient functioning of the thymus is critical for establishing and maintaining effective adaptive immunity^1^. Thymopoiesis is a highly complex process involving cross-talk between developing thymocytes and the non-hematopoietic supporting stromal microenvironment, primarily highly specialized thymic epithelial cells (TECs)^2^. However, the thymus is exceptionally sensitive to negative stimuli that, together with its well-characterized capacity for repair, leads to continual cycles of involution and regeneration in response to acute injury^3-5^. However, this capacity for regeneration declines with age as a function of thymic involution, which itself leads to a reduced capacity to respond to new pathogens, as well as poor response to vaccines and immunotherapy^6-9^. Therefore, there is a pressing clinical need for the development of therapeutic strategies that can enhance T cell reconstitution. One approach to therapeutic development is to exploit key factors that promote endogenous thymic regeneration into novel pharmacologic strategies to enhance T cell reconstitution in clinical settings of immune depletion such as HCT. However, the molecular mechanisms governing endogenous thymic regeneration are not fully understood.

Our recent studies have revealed that endogenous thymic repair is dependent on the production of two distinct regeneration factors, IL-23 and BMP4^10-12^. Both IL-23, by initiating the downstream production of IL-22 by innate lymphoid cells (ILCs)^12-15^, and BMP4 target TECs to facilitate regeneration^10-12^. These regeneration-associated factors have profound reparative effects in the thymus after acute injury; and can be utilized individually as therapeutic strategies of immune regeneration^5^. Although BMP4 and the IL-23-IL-22 axis represent two regenerative pathways that facilitate TEC repair; the damage-sensing mechanisms that trigger the production of these factors after damage from regeneration-initiating ECs and DCs are unknown.

## RESULTS

### NOD2 negatively regulates thymus regeneration by suppressing the production of BMP4 and IL-23

To identify potential mechanisms by which these regeneration networks are governed, we firstly performed transcriptome analysis on purified ECs (the main source of BMP4 after damage^10^) isolated from the mouse thymus at days 0 and 4 after a sublethal dose of total body irradiation (SL-TBI, 550cGy); timepoints that capture baseline levels and at the time of initiation of the regenerative response (day 4)^10,12^. Pathway analysis through the DAVID tool revealed a surprising number of pathways associated with immune function, which included most of the pathways with an FDR<0.05 (**Fig. 1a, S1a**). Further analysis identified that within these pathways, several genes shared a high frequency across all GO pathways (**Fig. 1b**). In particular, Nucleotide-binding oligomerization domain-containing protein 2 (NOD2) stood out not only as it is centrally involved in innate immunity via its role in detection of danger signals, such as bacterial peptidoglycan^16,17^, but also because NOD2 is one of the only molecular pathways found to suppress the production of IL-23 by DCs^18^. Therefore, we hypothesized that NOD2 activation was central to the inhibition of IL-23 and BMP4, and after damage this activation is abrogated upstream by the depletion of thymocytes. To explore the effects of NOD2 deficiency on thymic regeneration following insult, WT or *Nod2*^*-/-*^ animals were exposed to SL-TBI and levels of thymic IL-23 and BMP4 were measured by ELISA. *Nod2*^*-/-*^ thymi had increased absolute intracellular levels of IL-23 and BMP4 compared to WT controls (**Fig. 1c**). Consistent with this, there was a commensurate increase in thymic cellularity, from as early as 7 days, and even up to 18 weeks after SL-TBI (**Fig. 1d**), suggesting that these increased levels of regenerative factors are supporting superior thymic regeneration. Although there was no global change in thymocyte proportions (**Fig. 1e**), the enhanced total cellularity was reflected by increases in all subsets of thymocytes **(Fig. 1f)**. Remarkably, given their role in the production of the regenerative factors, there was no change in the number of ECs in the *Nod2*-deficient thymus; however, there was a greater regeneration of CD103+ DCs (**Fig. 1g**), which mediate IL-23 levels^12,19^. Importantly, given the increased levels of BMP4 and IL-23, which can directly or indirectly induce TEC proliferation and function^10,12^, there was significantly augmented regeneration of both cortical TECs (cTEC) and medullary TECs (mTECs) (**Fig. 1h**), both of which have distinct crucial roles in T cell development^2^. Notably, GSEA analysis supported the role of NOD2 in the damage response, with a significant enrichment in the signature of downstream NOD2 gene targets^20^ at day 4 after damage (**Fig. S1b**). Although the expression of *Nod2* in thymic tissue has been known for some time^21^, the only functional role for NOD2 that has been described in the thymus has been in thymocyte selection^22^.

**Figure 1:**
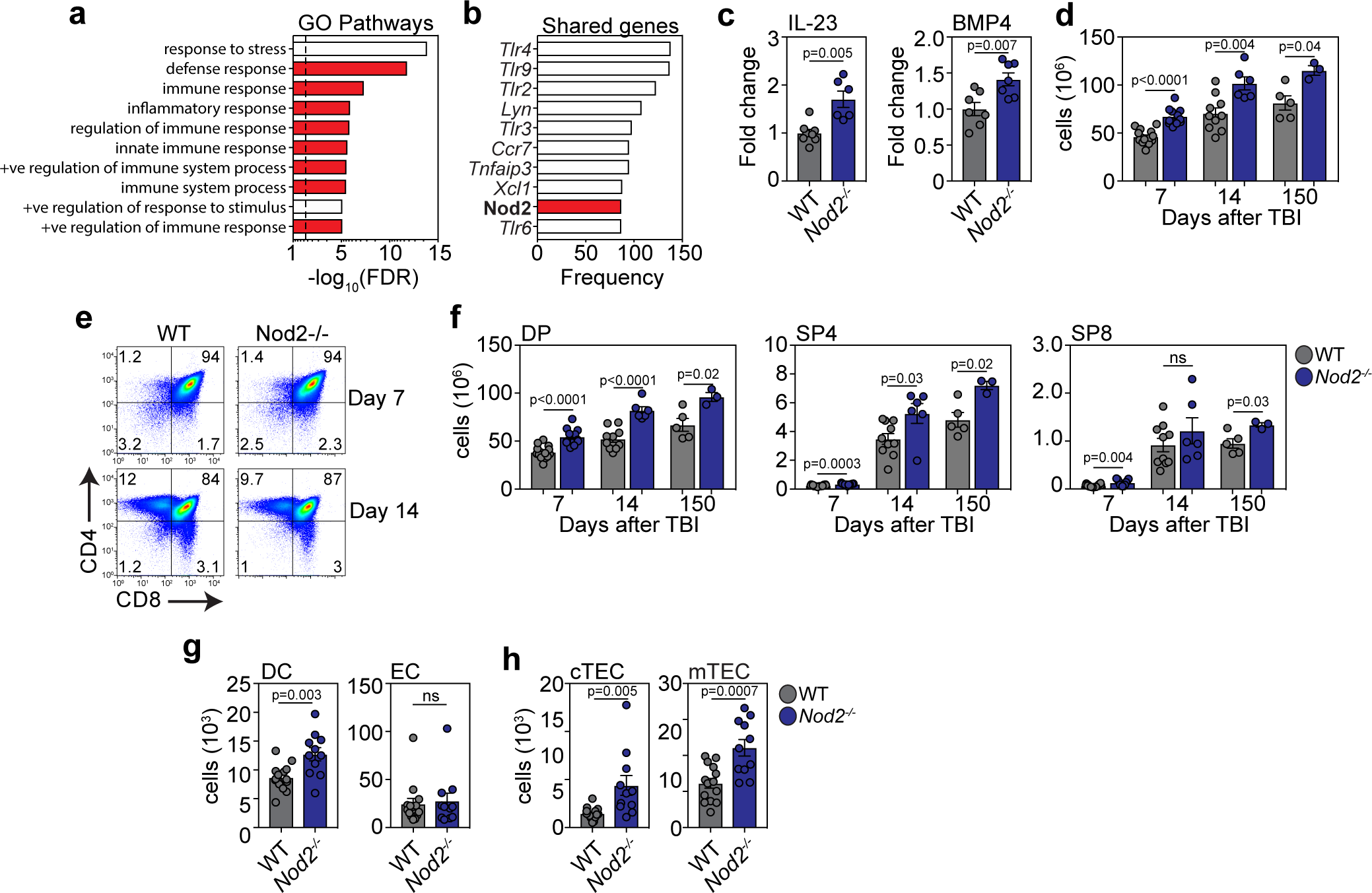
NOD2 limits thymus regeneration by inhibiting the production of regenerative factors. **a-b**, Thymuses were pooled from 6-week-old C57BL/6 mice and microarray analysis was performed on FACS purified ECs isolated from either untreated mice (d0) or 4 days after TBI (550 cGy, (d0, n=3; d4, n=2; each n pooled from 5 mice). **a**, Gene ontology (GO) pathway analysis was performed on upregulated genes at day 4 after SL-TBI using DAVID and the top ten pathways by FDR are displayed. Red bars represent pathways involved with immune function; dashed line at p=0.05. **b**, Top ten genes by frequency of representation amongst all GO pathways with an FDR <0.05. **c-h**, 6-8 week C57BL/6 WT or *Nod2*^*-/-*^ mice were given a sublethal dose of TBI (550 cGy) and thymus was harvested and analyzed at the indicated timepoints. **c**, Total thymic amounts of BMP4 and IL-23 were assessed at day 7 after SL-TBI (n=7/group across two independent experiments). **d**, Total thymic cellularity at days 7, 14, and 150 after SL-TBI (d7, n=11-15; d14, n=6-10; d150, n=3-5; all across two independent experiments except for d150). **e**, Concatenated flow plots showing CD4 and CD8 expression at days 7 and 14 after SL-TBI. **f**, Total number of CD4+CD8+ DP, CD4+CD3+ SP4, or CD3+CD8+ SP8 thymocytes (d7, n= 11-15; d14, n=6-10; d150, n=3-5; all across two independent experiments except for d150). **g**, Number of CD45+MHCII+CD11c+CD103+ DCs and CD45-EpCAM-CD31+PDGFRa-ECs at day 7 after SL-TBI (n=11-15/group across two independent experiments). **h**, Number of CD45^-^EpCAM^+^MHCII^+^UEA1^lo^Ly51^hi^ cTECs and CD45^-^EpCAM^+^MHCII^+^UEA1^hi^Ly51^lo^ mTECs at day 7 after SL-TBI (n=11-15/group across two independent experiments). Graphs represent mean ± SEM, each dot represents a biologically independent observation.

### miR29c mediates the suppressive function of NOD2

NOD2 has been previously shown to induce expression of the microRNA miR29 by directly regulating the transcription of *Il12p40*, and indirectly regulating *Il23p19*^18^. MicroRNAs are small non-coding RNAs that critically govern protein expression by binding to and degrading target RNA^23-25^. There are three members of the miR29 family, each with some overlapping physiological functions; including a TEC-intrinsic role of miR29a in regulating the response to interferon signaling and Aire-dependent gene expression^26,27^. All three miR29 family members increase in the aged thymus, suggesting a potential role in involution^28^. To understand if miR29 was involved in the regenerative response in the thymus, we first assessed the expression levels of mature 3p and 5p arms of miR29a, miR29b, and miR29c in the thymus from WT or *Nod2*^*-/-*^ mice 3 days after TBI and found lower expression of miR29c-5p in *Nod2*-deficient mice **(Fig. 2a)**. To elucidate if there was a cell-specific function of miR29 in the thymus we assessed miR29 expression in purified populations of ECs, DCs and DP thymocytes, and did not find a significant change in the expression of either miR29a-5p or miR29b-5p after damage (Fig. **S2a**). However, we observed decreased expression of miR29c-5p in ECs and DCs after damage, with this damage-induced reduction lacking in DP thymocytes (**Fig. 2b**), suggesting an endogenous regulation in miR29c-5p after damage in the regeneration-initiating DCs and ECs; both of which have higher miR29c-5p expression at baseline compared to DP thymocytes (**Fig. S2b**).

**Figure 2:**
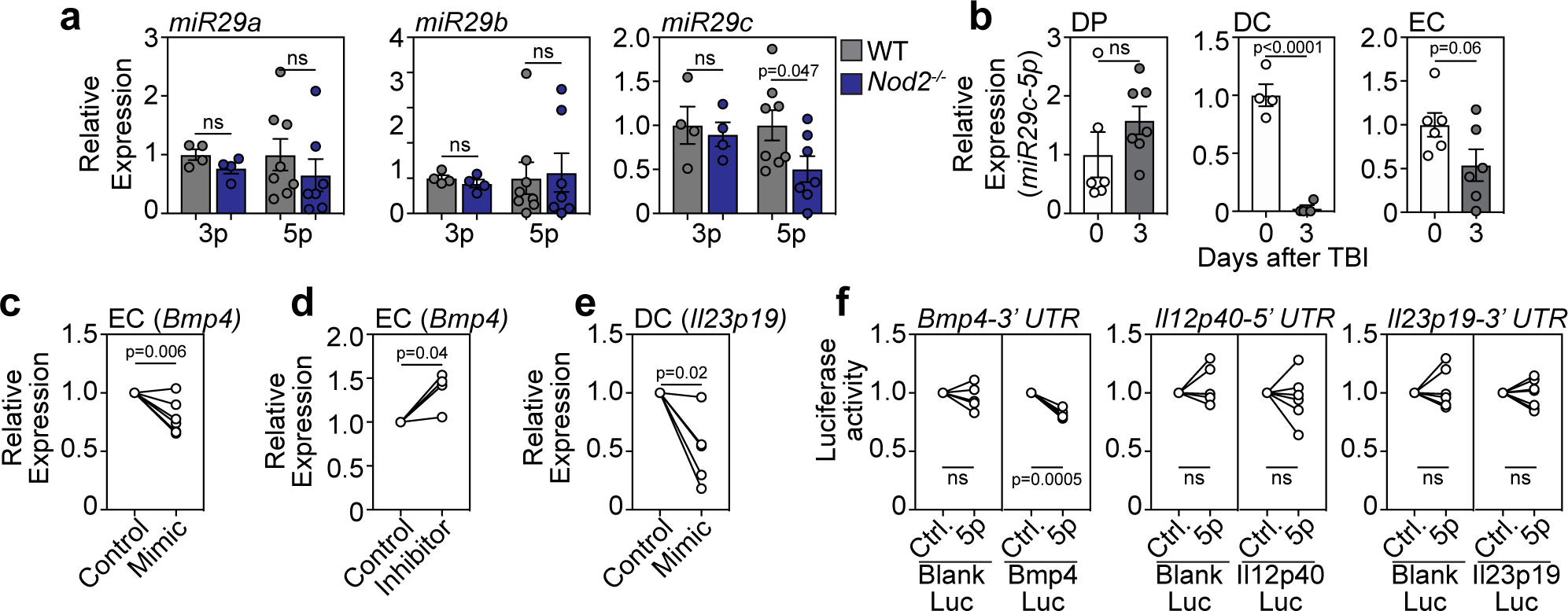
miR29c mediates the effects of NOD2 in limiting production of regenerative factors in ECs and DCs. **a**, Thymuses were isolated from 6-8 week C57BL/6 WT or Nod2-/- mice at day 3 following SL-TBI and expression of 3p and 5p arms of miR29a, miR29b, or miR29c was analyzed by qPCR (3p, n=4/group; 5p, n=7/group across two independent experiments). **b**, DPs, ECs and DCs were FACS purified from WT thymuses at day 0 or 3 after SL-TBI and expression of miR29c-5p was analyzed by qPCR (DCs, n=4; ECs/DPs, n=6-7/population/timepoint across two independent experiments). **c**, exECs were generated as previously described11,31 and transfected with a miR29c mimic. 20 hours after transfection, expression of Bmp4 was analyzed by qPCR (n=7 independent experiments). **d**, exECs were transfected with a miR29c inhibitor and expression of Bmp4 was analyzed by qPCR 20 hours after transfection (n=4 independent experiments). **e**, CD11c+ DCs were isolated by magnetic enrichment from untreated C57BL/6 thymus and transfected with a miR29c mimic. 20 hours after transfection, Il23p19 was analyzed by qPCR (n=5 across three independent experiments). **f**, HEK293 cells were co-transfected with either BMP4-3’UTR, Il12p40-5’UTR, or Il23p19-3’UTR luciferase constructs and a miR29c-5p mimic. Binding activity was quantified by measuring luciferase activity after 20 hours (n=5-7 independent experiments). Graphs represent mean ± SEM, each dot represents a biologically independent observation.

To functionally assess the relationship between miR29c-5p and the production of regenerative factors, we used a technique to constitutively activate the Akt pathway in freshly isolated thymic ECs using the pro-survival adenoviral gene *E4ORF1*, which allows for their *ex vivo* propagation and expansion while maintaining endothelial phenotype, adaptability, as well as vascular tube formation capacity^10,29^, allowing for functional manipulation of the cells for *in vitro* modeling of regenerative pathways. Consistent with the decrease of miR29c-5p in regeneration-initiating cells after damage, and the previously identified role of miR29 in the attenuation of *Il23p19*^*18*^, we hypothesized that miR29c-5p regulates the expression of *Bmp4* in ECs. Overexpression of miR29c-5p in thymic exECs resulted in a significant reduction in the expression of *Bmp4* (**Fig. 2c**), which was reversed upon inhibition of miR29c-5p **(Fig. 2d)**. Freshly isolated DCs transfected with a miR29c-5p mimic demonstrated a similar reduction in expression of the *Il23* p19 subunit (**Fig. 2e**), suggesting that miR29c-5p negatively regulates these regenerative factors in both ECs and DCs. miRNAs regulate gene expression in several ways; either by binding to the 3’ UTR of their target transcript and inducing post-transcriptional modifications and translational repression, ultimately resulting in transcript degradation; or by binding to the 5’ UTR sequence of their target transcript and exerting silencing effects on gene expression^30^. To understand if miR29c-5p regulation of *Bmp4* and *Il23p19* was due to direct binding, and to determine where this binding occurred, we carried out a luciferase assay using 3’UTR- and 5’UTR BMP4-luciferase constructs that were co-transfected with a miR29c-5p mimic. We observed a degradation of the 5’UTR of *Bmp4* in the presence of the miR29c-5p mimic, identifying a direct and specific regulation **(Fig. 2f)**, with a suggested indirect effect of miR29c-5p on *Il23p19* stability, consistent with previous reports^27^.

### Detection of apoptotic thymocytes by TAM receptors suppresses the production of regenerative factors

Although we have demonstrated that attenuation of NOD2-dependent miR29c-5p expression is involved in executing the regenerative response after damage by regulating levels of critical regenerative factors, the upstream initiator of this cascade is not defined. In our previous studies, we found a link between the loss of thymic cellularity and the initiation of the IL-22 and BMP4 pathways^10-12^. In the case of IL-22/IL-23, this could be directly correlated with the depletion in the number of CD4^+^CD8^+^ double positive (DP) thymocytes, as mice with a genetic block before the DP stage (*Rag1*^*-/-*^, *Il7ra*^*-/-*^, *Il7*^*-/-*^ *and Tcrb*^*-/-*^)^31^, or mice treated with dexamethasone which causes a targeted reduction in DP thymocytes^32^, expressed profoundly more IL-22 and IL-23 than WT controls or mutant mice with blocks further downstream in T cell development (*Tcra*^*-/-*^ and *Ccr7*^*-/-*^)^12^. In the case of the mutant mouse strains, this occurred even without acute thymic damage, suggesting that merely the absence of DP thymocytes is enough to trigger these reparative pathways. Although 80-90% of the thymus is comprised of DP thymocytes, approximately 99% of these cells undergo apoptosis under homeostatic conditions due to selection events^33,34^. Taken together with reports that apoptotic cells can regulate the production of cytokines, including reducing the production of IL-23 by DCs^35^, we hypothesized that the presence of homeostatic apoptotic thymocytes can limit the production of the regenerative factors IL-23 and BMP4. To test this, we performed co-culture experiments of thymocytes with thymic ECs or DCs, where the thymocytes had been induced to undergo apoptosis with dexamethasone^36^ (referred to as apoptotic cells, ACs), or where apoptosis was inhibited with the pan-caspase inhibitor z-VAD-FMK. Consistent with the hypothesis that ACs are suppressive to the expression of *Bmp4* and levels of IL-23, when apoptosis was inhibited in thymocytes there was a significant increase in the expression of both of *Bmp4* in ECs and IL-23p19 in DCs (**Fig. 3a, b**). Indeed, Annexin V binding was reduced in thymocytes incubated with z-VAD-FMK, indicating a reduction in apoptotic thymocytes (**Fig. S3**). Annexin V is a surrogate marker of apoptotic cells and binds to exposed phosphatidylserine (PtdSer); inversion of which from the inner cell membrane is a key identifying feature of apoptotic cells^37,38^. Even though on a per cell basis there is an increase in Annexin V binding on DP thymocytes after damage (**Fig. 3c**), given the severe depletion in DP thymocytes after damage (**Fig. 3d, e**), there was an overall profound decrease in the total exposed PtdSer (**Fig. 3f, S4**), resulting in a robust correlation between exposed PtdSer and DP cellularity (**Fig. 3g**). Apoptotic cell clearance is facilitated by the detection of PtdSer on the apoptotic cells by TAM receptors, Tyro, Axl, and Mer, on nearby cells. TAM receptors are a family of transmembrane tyrosine kinase receptors that recognize their extracellular ligands Gas6 or ProS 1 in the presence of PtdSer, and have a critical role in the maintenance of immune homeostasis, including regulation of thymic negative selection^39-41^. Both thymic ECs and DCs express *Axl, Mer* and *Tyro3* (**Fig. 3h)**, as has been extensively reported previously in other tissues^40^, and upon inhibition of TAM receptors with the pan-inhibitor RXDX-106^42^, we found increased expression of *Bmp4* in ECs and IL-23 in DCs in the presence of ACs, suggesting that TAM receptors mediate suppressive signals from ACs (**Fig. 3i, j**).

**Figure 3:**
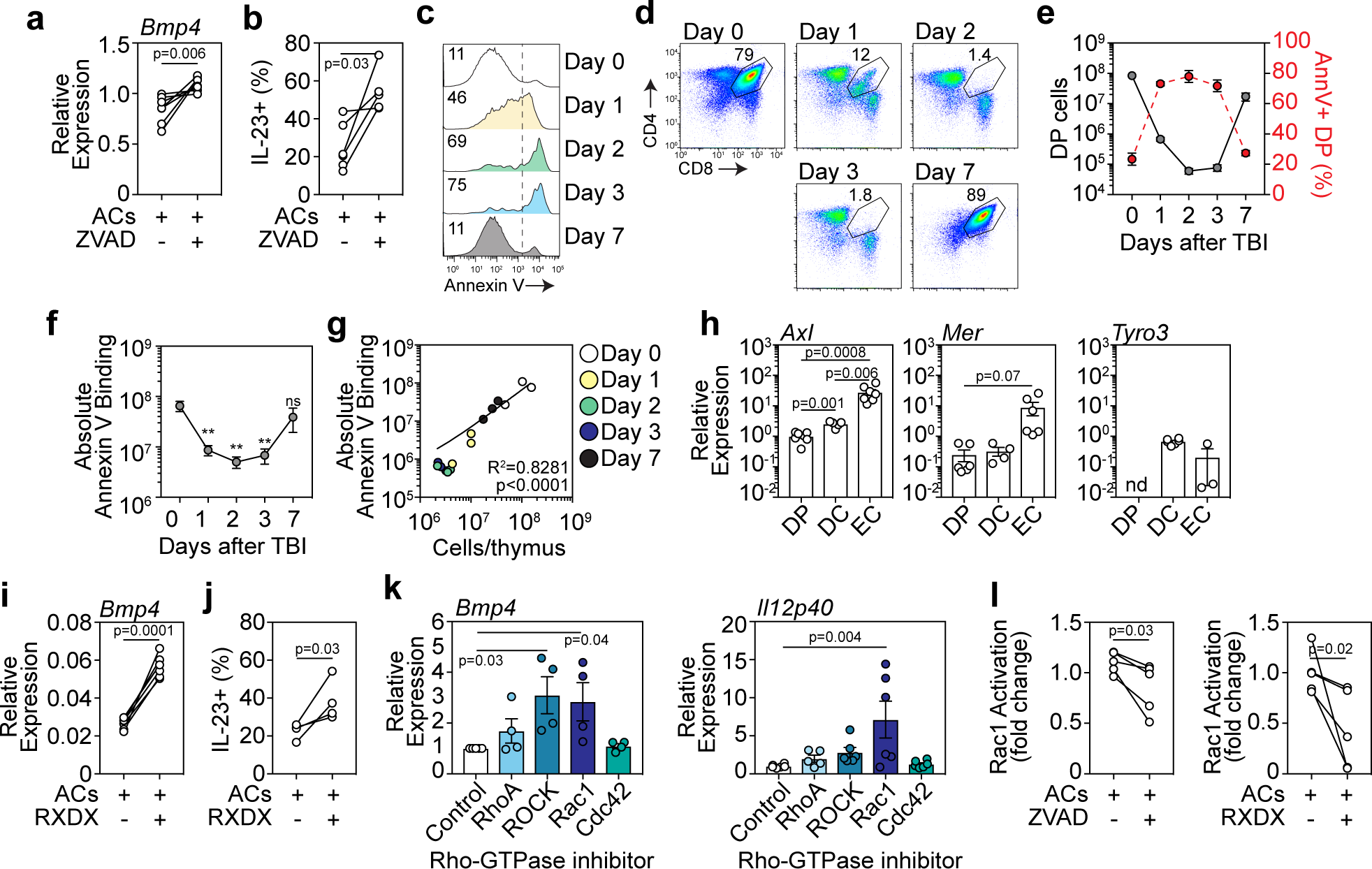
TAM receptor detection of phosphatidylserine mediates thymocyte suppression of regenerative factors. **a-b**, Thymocytes were isolated from untreated C57BL/6 mice and incubated for 4 hours with Dexamethasone (100 nM) or zVAD-FMK (20 µM). After 4 hours, apoptotic thymocytes (ACs) were washed and co-cultured with exECs **(a)** or freshly isolated CD11c+ DCs **(b)** for 24 hours after which Bmp4 expression was analyzed by qPCR (n=8 across 5 independent experiments) or IL-23 was analyzed by intracellular cytokine staining (n=5-6 across two independent experiments). **c-g**, 6-8 week old C57BL/6 mice were given sublethal TBI (550 cGy) and thymus harvested at days 0, 1, 2, 3, and 7. **c**, Annexin V staining on CD4+CD8+ DP thymocytes (displayed are concatenated plots from 3 individual mice, representative of three independent experiments). **d**, Staining for CD4 and CD8 on thymus cells (displayed are concatenated plots from 3 individual mice, representative of three independent experiments). **e**, Total number of DP thymocytes (solid line; left axis) compared with proportion of Annexin V+ DP thymocytes (red broken line; right axis) (n=8 across 3 independent experiments). **f**, Absolute binding of Annexin V in the thymus (n=8 across 3 independent experiments). **g**, Correlation of absolute thymic binding of Annexin V with total number of cells in the thymus (n=3/timepoint comprising one of three independent experiments). **h**, DP thymocytes (CD45+CD4+CD8+), DCs (CD45+CD11c+MHCII+), or ECs (CD45-EpCAM-CD31+) were FACS purified from untreated C57BL/6 mice and expression of Axl, Mer, and Tyro3 were analyzed by qPCR (DP, n=7; DC, n=5; EC, n=7 across two independent experiments). **i-j**, Thymocytes were isolated from untreated C57BL/6 mice and incubated for 4 hours with dexamethasone (100 nM). After 4 hours, apoptotic thymocytes (ACs) were washed and co-cultured with exECs **(i)** or freshly isolated CD11c+ DCs **(j)** in the presence or absence of the TAM receptor inhibitor RXDX-106 (25μM) for 20 hours after which Bmp4 expression was analyzed by qPCR (n=6/treatment across 2 independent experiments) or IL-23 was analyzed by intracellular cytokine staining (n=4). **k**, exECs or freshly isolated CD11c+ DCs were incubated for 20 hours in the presence of inhibitors (all at 50μM) for RhoA (Rhosin), ROCK (TC-S 7001), Rac1 (EHT-1864), or Cdc42 (ZCL272) after which Bmp4 (ECs) or Il12p40 were analyzed by qPCR (exECs, n=5 independent experiments; DCs, n=6 across two independent experiments). **l**, Thymocytes were isolated from untreated C57BL/6 mice and incubated for 4 hours with dexamethasone (100 nM) or ZVAD-FMK (20μM). After 4 hours, apoptotic thymocytes (ACs) were washed and co-cultured with exECs in the presence or absence of RXDX-106 (25μM) for 20 hours after which Rac1-GTPase activation was measured using a GTPase ELISA specific for Rac1 (n=5 across two independent experiments). Graphs represent mean ± SEM, each dot represents a biologically independent observation.

### TAM receptor signaling promotes Rac1 GTPase activation and limits *Bmp4* and *Il23*

PtdSer guides and strengthens ligand binding to TAM receptors and is critical for TAM receptor activation and downstream signaling^40^. One of the common downstream targets of TAM receptors are Rho GTPases, which is notable given recent reports that, in addition to its capacity to sense bacterial-derived peptidoglycans, NOD2 can also act as a cytosolic sensor of activated Rho GTPases^43,44^. Rho GTPases have multiple cellular roles centered on modulation of the cellular cytoskeletal architecture, and regulate processes such as cell adhesion, migration, polarization and trafficking^45^. In the thymus, Rac1, RhoA and Cdc42 have all been implicated in aspects of β-selection and positive selection^46-48^, two stages of T cell development with high levels of thymocyte apoptosis^34^. This strengthens the putative link between loss of PtdSer-exposed cells in the thymus (and the loss of downstream Rho GTPase activation) triggering the production of regenerative factors after damage. To investigate this relationship, we first performed a functional screen of inhibitors of Rho, Rac, and Cdc42 subclasses of Rho GTPases^45,49,50^ in exECs or freshly isolated thymic DCs and assessed the expression of regenerative factors. Using this approach, we identified that while inhibition of Cdc42 did not mediate any effect on the production of *Bmp4* by ECs or *Il12p40* in DCs, inhibition of RhoA and Rac1 led to significantly increased expression in these regenerative factors (**Fig. 3k**). Given the common profound effect of Rac1 inhibition across ECs and DCs in these *in vitro* assays, we next sought to determine if detection of apoptotic thymocytes could indeed lead to Rac1 GTPase activation. Consistent with our proposed framework, we demonstrated that apoptotic thymocytes induced activation of Rac1 in thymic ECs, an effect that was reversed when co-cultured with apoptosis-inhibited thymocytes or by inhibition of TAM receptors with RXDX-106 **(Fig 3l**).

### Rac1 GTPase inhibition enhances thymus regeneration and thymic output *in vivo*

Several strategies targeting Rho GTPases have been examined pre-clinically for multiple cancers. These include small molecules targeting the spatial regulation of GTPase activators^51^, and Rho-GEF interactions, such as Rhosin^52^, and our therapeutic candidate Rac1 GTPase inhibitor EHT1864^53^, with the greatest clinical trial success thus far in targeting the downstream kinase of RhoA, ROCK^54,55^. To determine if this pathway could be manipulated for therapeutic efficacy in thymic regeneration, we treated mice with the Rac1 GTPase inhibitor EHT1864 following SL-TBI and identified a robust regeneration of thymic cellularity (**Fig. 4a**). EHT1864 had no effect on thymus cellularity in *Nod2*^*-/-*^ mice (**Fig. 4a**), supporting our hypothesis of a Rac1-NOD2 regeneration axis. Although thymocyte proportions were unaffected by treatment with EHT1864 (**Fig. 4b**), we found a similar increase in almost all thymocyte subsets (**Fig. 4c**), which was not observed in EHT1864 treated *Nod2*-deficient mice (**Fig. S5**). Moreover, a significant decrease in miR29c-5p expression was determined (**Fig. 4d**), with an increase in the levels of BMP4 and IL-23 (**Fig. 4e**), accompanied by the enhanced TEC regeneration after EHT1864 treatment (**Fig. 4f**). In the periphery, while there was no change in the number of CD4+ or CD8+ thymocytes or their relative proportions (**Fig. 4g, h**), there was an increase in the output of naïve T cells in mice treated with EHT1864, reflected by their proportion and the ratio of naïve:memory cells (**Fig. 4i, j**).

**Figure 4:**
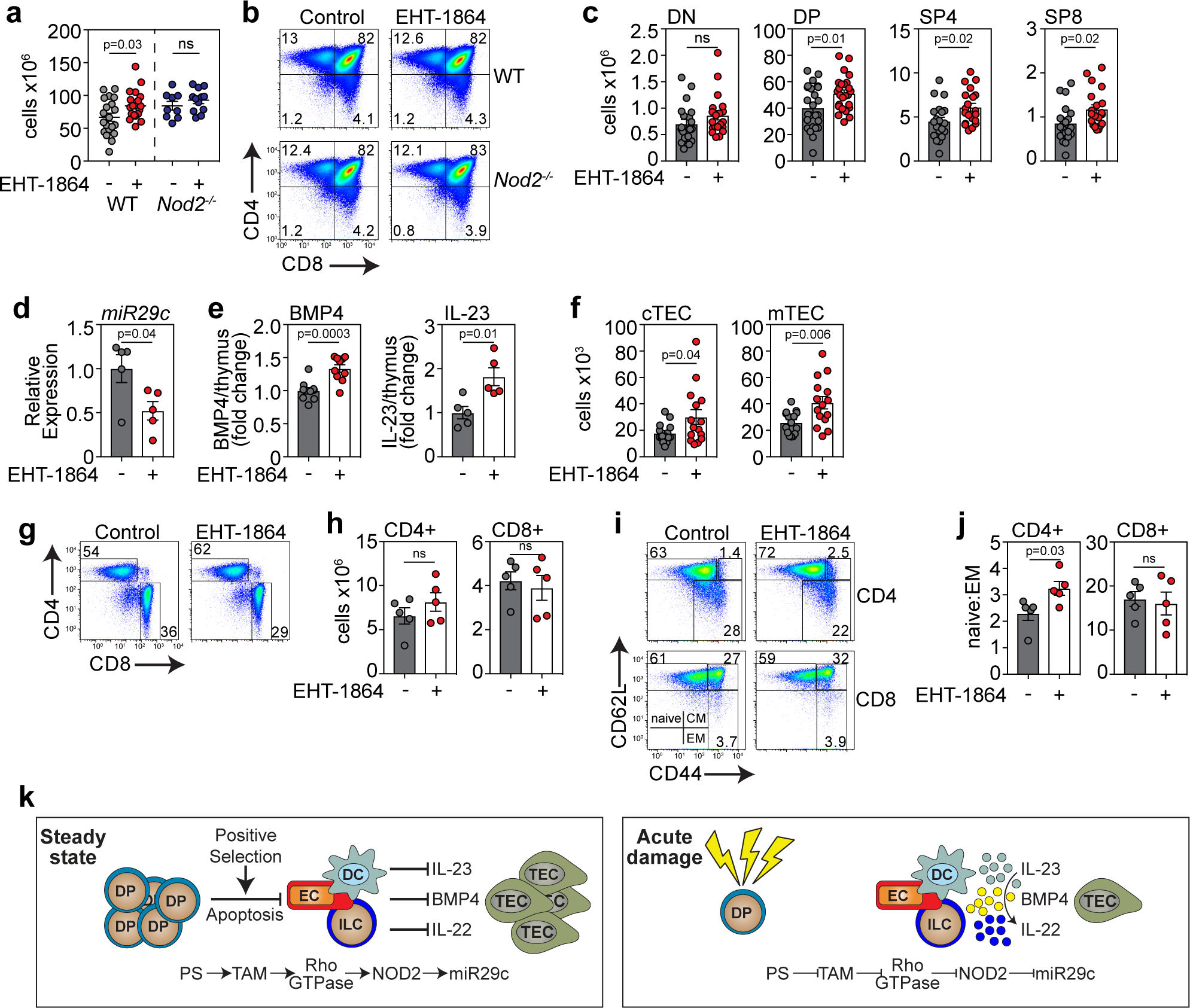
Rac1 inhibition enhances thymus regeneration and peripheral CD4+ naïve T cell recovery after acute damage. 6-8 week C57BL/6 WT or Nod2-/- mice were treated with the Rac1 inhibitor EHT1864 (40 mg/kg i.p. injection) at days 3, 5 and 7 following a sublethal dose of TBI (550 cGy). **a**, Total thymic cellularity at day 14 after SL-TBI (WT, n=20/treatment across 4 independent experiments; KO, n=9-12 across three independent experiments). **b**, Flow plots showing CD4 and CD8 expression at day 14 after SL-TBI (gated on CD45^+^ cells; plots concatenated of all samples in each treatment group from one experiment). **c**, Total number of CD4^+^CD8^+^ DP, CD4^+^CD3^+^ SP4, or CD3^+^CD8^+^ SP8 thymocytes (WT, n=20/treatment across 4 independent experiments; KO, n=9-12 across three independent experiments). **d**, Expression of miR29c analyzed by qPCR (n=5/group). **e**, Total thymic amounts of BMP4 and IL-23 assessed by ELISA (BMP4, n=10/group across two independent experiments; IL-23, n=5). **f**, Total number of CD45^-^EpCAM^+^MHCII^+^UEA1^lo^Ly51^hi^ cTECs and CD45^-^EpCAM^+^MHCII^+^UEA1^hi^Ly51^lo^ mTECs (n=15/group across three independent experiments). **g**, Flow plots showing CD4 and CD8 expression in the spleen at day 56 after SL-TBI (gated on CD45+CD3+ cells). **h**, Total number of CD4+ and CD8+ T cells in the spleen at d56 (n=5/group). **i**, Plots of CD62L and CD44 on CD4^+^ and CD8^+^ T cells (gated on either CD45^+^CD3^+^CD4^+^CD8^-^ or CD45^+^CD3^+^CD4^-^CD8^+^) cells; plots concatenated of all samples in a given experiment). **j**, Ratio of number of naïve (CD62L^hi^CD44^lo^) CD4^+^ or CD8^+^ to CD4^+^ or CD8^+^ EM (CD62L^hi^CD44^hi^) T cells (n=5/group). Graphs represent mean ± SEM.

## DISCUSSION

Despite its importance for generating a competent repertoire of T cells, the thymus is exquisitely sensitive to acute injury such as that caused by infection, shock, or common cancer therapies such as cytoreductive chemo- or radiation therapy; however, also has a remarkable capacity for repair^4,12,56^. Even in the clinical setting where children who have had large parts of their thymus removed exhibit significant thymic repair^4^. Thus, endogenous thymic regeneration is a critical process to restore immune competence following thymic injury. Although there is likely continual thymic involution and regeneration in response to stress and infection in otherwise healthy people, acute and profound thymic damage such as that caused by common cancer cytoreductive therapies, conditioning regimes as part of hematopoietic cell transplantation (HCT), or age-related thymic involution, leads to prolonged T cell deficiency; precipitating high morbidity and mortality from opportunistic infections and may even facilitate cancer relapse^8,9,57-66^. Of note, the general phenomena of endogenous thymic regeneration has been known for longer even than its immunological function^67,68^, however, the underlying mechanisms controlling this process have been largely unstudied^6,69^.

Our recent studies have revealed that thymic ILCs and ECs, through their production of the regeneration-associated factors IL-22 and BMP4, respectively, have profound reparative effects in the thymus after acute injury^10-12^, and both pathways act by stimulating TECs^2^. Our findings suggest this high homeostatic level of apoptosis; signaling through TAM receptors and the downstream activation of Rac1 GTPase, NOD2, and miR29c; suppresses the steady-state production of the regenerative factors BMP4 ad IL-23 in in ECs and DCs, respectively. However, after damage, the loss of thymocytes (and their exposed PtdSer) abrogates this tonic signaling, alleviating this suppression and inducing production of BMP4 and IL-23. Given the passive activation of this pathway, this acts as a “dead-man’s switch” that ensures faithful regulation and activation of the regeneration program in settings of damage.

Notably, while expression of *Nod2* in the thymus has been known for some time^21^, it has thus far only been functionally linked to positive selection and maturation of CD8+ T cells by facilitating TCR-ERK signaling^70^. It has been previously reported that NOD2 controls the production of IL-23 by DCs via miR29, which directly regulated the transcription of *Il12p40* and indirectly through *Il23p19*^*18*^. This is consistent with the finding that NOD2 can negatively influence the production of IL-12, but not other cytokines such as IL-10 and TNF^71^. Although the three members of the miR29 family have overlapping physiological roles^27,72-78^, we found that miR29c-5p facilitates NOD2-mediated suppression of thymic regeneration directly by suppressing *Bmp4* expression, and indirectly regulating both p19 and p40 subunits of IL-23. In the thymus we can detect miR29a, miR29b and miR29c in DP thymocytes, ECs and DCs, although expression of miR29c-5p is significantly higher in ECs and DCs than DP thymocytes. This differential expression is particularly interesting in light of our finding that after SL-TBI there is a significant loss of miR29c-5p in ECs and DCs, but not DP thymocytes, and significantly less expression of miR29c-5p in *Nod2*^*-/-*^ compared to WT controls after damage. It is also notable that both NOD2-miR29 and apoptotic cells have been identified to negatively regulate the production of IL-23 by DCs^18,79^. Furthermore, there is evidence that miR29 can modulate thymic involution in response to infection via regulation of the interferon-*α* receptor^27^.

The canonical ligands for NOD2 are peptidoglycans found in the cell wall of bacteria^16,80^; however, these are unlikely to serve as a NOD2 activator in the thymus since it is typically thought of as a sterile organ^81^. One recently described alternate function of NOD2 is as a cytosolic sensor of activated Rho GTPases^43,44,82-85^. The Rho GTPase family is responsible for a wide range of physiological processes^49,86,87^, including the intrathymic regulation by Rac1, RhoA and Cdc42 of β-selection and positive selection^47,48^. Rho GTPases including RhoA, Rac1 and Cdc42 can be activated by caspase-3 during apoptosis^88-90^, and themselves further promote apoptosis^91-93^, including in thymocytes that do not successfully express a pre-TCR ^94^. Taken together, these findings suggest that that Rho GTPases could act as a mediator of NOD2 signaling during steady state thymopoiesis.

Although endogenous thymic regeneration is required for renewal of immune competence following everyday insults, regeneration can be a prolonged process representing an important clinical problem in older patients. Age-related thymic deterioration leads to a decline in the output of newly generated naïve T cells from the thymus, subsequent constriction in the breadth of the peripheral TCR repertoire, coupled with the compensatory expansion of existing naïve and memory T cells, and ultimately culminates in suppressed immunogenicity to new antigens ^95^, whether they be new infections, vaccinations, or cancer neoantigens. Therefore, effective therapeutic targeting of these endogenous pathways is crucial. Several strategies targeting Rho GTPases have been examined pre-clinically for multiple cancers. These include small molecules targeting the spatial regulation of GTPase activators ^51^, and Rho-GEF interactions, such as Rhosin ^52^ and our therapeutic candidate EHT1864 ^53^. Although several Rho GTPases, including Rac1, RhoA and Cdc42 play a role in thymopoiesis^47,48^, notably during stages of selection when there is with high levels of apoptosis^34,96^; however, given the therapeutic window when thymocytes are largely absent, and the redundancy in Rac1 and Rac2 in this process^97^, suggests that Rac1 inhibition represents a promising therapeutic strategy for boosting immune function.

These findings not only uncover mechanisms governing endogenous thymic repair; they outline a previously unappreciated master regulatory mechanism of regeneration that may be applicable in other high turnover tissues such as gut, liver, kidney, and skin. Furthermore, by targeting this pathway pharmacologically, we propose a novel therapeutic intervention that could be used to boost immune function in patients whose thymus has been damaged due to age, cytoreductive therapies, infection or other causes.

## METHODS

### Mice

Inbred male and female C57BL/6 mice were obtained from the Jackson Laboratories (Bar Harbor, USA). *Nod2*^*-/-*^ (B6.129S1-*Nod2*^*tm1Flv*^/J) mice were obtained from Jackson Laboratories and bred in house. All experimental mice were used between 6-8 weeks old. To induce thymic damage, mice were given sub-lethal total body irradiation (SL-TBI) at a dose of 550 cGy from a cesium source mouse irradiator (Mark I series 30JL Shepherd irradiator) with no hematopoietic rescue. For *in vivo* studies of EHT-1864 administration, mice were given SL-TBI (550cGy) and subsequently received i.p. injections of 40 mg/kg EHT1864 (3872, Tocris, UK), or 1 x PBS as control, on days 3, 5 and 7 following TBI. Mice were maintained at the Fred Hutchinson Cancer Research Center (Seattle, WA), and acclimatized for at least 2 days before experimentation, which was performed per Institutional Animal Care and Use Committee guidelines.

### Cell Isolation

Single cell suspensions of freshly dissected thymuses were obtained and either mechanically suspended or enzymatically digested as previously described^12,98^ and counted using the Z2 Coulter Particle and Size Analyzer (Beckman Coulter, USA). For studies sorting rare populations of cells in the thymus, multiple identically-treated thymuses were pooled so that sufficient number of cells could be isolated; however, in this instance separate pools of cells were established to maintain individual samples as biological replicates.

### Reagents

Cells were stained with the following antibodies for analysis CD3-FITC (35-0031, Tonbo Bioscience), CD8-BV711 (100748, BioLegend), CD4-BV650 (100546, BioLegend), CD45-BUV395 (565967, BD Biosciences), CD90-BV785 (105331, BioLegend), CD11c-APC (20-0114, Tonbo Biosciences), MHC-II-Pac Blue (107620, BioLegend), CD103-PercPCy5.5 (121416, BioLegend), CD11b-A700 (557960, BD Pharmingen), EpCAM-PercPe710 (46-5791-82, eBioscience), Ly51-PE (12-5891-83, eBioscience), CD31-PECy7 (25-0311-82, eBioscience), CD140a-APC (135907, BioLegend), UEA1-FITC (FL-1061, Vector Laboratories), TCRbeta-PECy7 (109222, BioLegend), CD62L-APC-Cy7 (104427, BioLegend), CD44-Alexa Fluor RTM700 (56-0441-82, BioLegend), CD25-PercP-Cy5.5 (102030, BioLegend). Annexin V staining (640906, BioLegend) was performed in Annexin V binding buffer (422201, BioLegend). Flow cytometric analysis was performed on an LSRFortessa X50 (BD Biosciences) and cells were sorted on an Aria II (BD Biosciences) using FACSDiva (BD Biosciences) or FlowJo (Treestar Software).

### Generation of exECs

exECs were generated as previously described ^29^. Briefly, CD45^-^CD31^+^ cells were FACS purified and incubated with lentivirus containing either the E4ORF1 or myrAkt construct for 48 hours. Cell culture medium containing 20 % FBS (SH30066.03, HyClone, GE Life Sciences), 10 mM HEPES (15630-080, Invitrogen), 1 % Glutamax (35050061, Life Technologies), 1 % Non-Essential Amino Acids (11140050, Life Technologies), 1% PenStrep (15240-062, Invitrogen), 50 ug/ml Heparin (H3149, Sigma), 50 ug/ml Endothelial Cell Supplement (02-102, Millipore-Sigma), 5 µM SB431542 (1614/10, R&D Systems), 20 ng/ml FGF (100-18B, Peprotech) and 10 ng/ml VEGF (450-32, Peprotech) at 37 °C, 5 % O_2_, 5 % CO_2_ in a HERAcell 150i incubator (Thermo Fisher, USA).

### Co-culture experiments

Co-culture experiments were carried out using exECs or DCs and thymocytes harvested from mechanically dissociated thymus from untreated mice, or in the case of DC analysis whole thymus cultures were used. Harvested thymocytes were incubated with either 100 nM dexamethasone (D2915, Sigma Aldrich, Germany), or 20 µM z-VAD-FMK (2163, Tocris, UK) for 4 hours at 37 °C prior to co-culture, washed twice with PBS, and resuspended in exEC media for co-culture (1 ⨯ 10^6^ cells / well). Cells were harvested 20 hours post co-culture and prepared for either qPCR analysis or flow cytometry analysis. Thymic DCs were isolated from untreated mice using CD11c UltraPure microbeads (130-108-338, Miltenyi Biotech, USA), on enzymatically digested thymuses. DCs were cultured in DMEM (11965, Gibco), 10% FBS (SH30066.03, HyClone, GE Life Sciences), and 1% PenStrep (15240-062, Invitrogen). For TAM receptor inhibitor studies, exECs were treated with 25 µM RXDX-106 (CEP-40783, s8570, Selleck Chemicals) 30 minutes prior to incubation with dexamethasone treated or z-VAD-FMK treated thymocytes, and *Bmp4* expression was determined by qPCR analysis 20 h post co-culture.

### ELISA

Thymuses were homogenized in RIPA buffer (25 mM Tris pH 7.6, 150 mM NaCl, 1% NP-40, 0.1% SDS, 0.05% sodium deoxycholate, 0.5 mM EDTA) with protease inhibitors (Thermo, A32955), using a Homogenizer 150 (Fisher Scientific) at a concentration of 10 mg/ml, where protein concentration was further normalized using BCA assay. BMP4 (DY485-05, R&D Systems) and IL-23 (433704, BioLegend) levels were assessed by ELISA, and absorbance was measured on the Tecan Spark 10M (Tecan, Switzerland).

### Microarray

Microarray analysis was performed on an Affymetrix MOE 430 A 2.0 platform in triplicate for untreated mice as well as day 4 after TBI. To obtain sufficient RNA for every timepoint, thymic ECs of several mice were pooled. All samples underwent a quality control on a bioanalyzer to exclude degradation of RNA. GSEA analysis was performed using the GSEA tool v4.1 of the Broad Institute (http://software.broadinstitute.org/gsea). Comparisons were made to known signaling pathways from the Gene Expression Omnibus (GEO) database (GSE226611). Pathway analysis was performed by submitting genes changed >1.5 (p<0.05) to DAVID Bioinformatics Resource v6.8^99,100^. Microarray data generated from ECs at day 4 after TBI will be deposited in the GEO; but was generated concurrently to untreated day 0 EC data, which has already been deposited to the GEO under number GSE106982^10^.

### qPCR

RNA was extracted from exECs or DCs using a RNeasy Mini kit (74104, Qiagen), and from sorted cells using a RNeasy Plus Micro kit (*74034, Qiagen*). cDNA was synthesized using the iScript gDNA Clear cDNA Synthesis kit (1725035, Bio-Rad, USA) and a Bio-Rad C1000 Touch ThermoCycler (Bio-Rad). RNA expression was assessed in the Bio-Rad CFX96 Real Time System (Bio-Rad), using iTaq Universal SYBR Green Supermix (1725122, Bio-Rad), and the following primers: *B-Actin* (*F 5’-CACTGTCGAGTCGCGTCC-3’; R 5’-TCATCCATGGCGAACTGGTG-3’*); *Il12p40* (*F 5’-AAGGAACAGTGGGTGTCCAG-3’, R 5’-CATCTTCTTCAGGCGTGTCA-3’*); *Il23p19* (*F 5’-GACTCAGCCAACTCCTCCAG-3’; R 5’-GGCACTAAGGGCTCAGTCAG-3’*); *Bmp4* (*qMmuCED0046239, Bio-Rad*). miRNA was extracted from cells using an miRNeasy Mini kit (217004, Qiagen) or miRNeasy Micro kit (1071023, Qiagen), and cDNA was synthesized using a Taqman Advanced miRNA cDNA Synthesis kit (A28007, Thermo Fisher). miRNA expression was measured on a Bio-Rad CFX96 Real Time System (Bio-Rad), using Taqman Advanced Master Mix (4444557, Thermo Fisher) and the following primers (Thermo Fisher): miR29a-3p (mmu478587_mir); miR29b-3p (mmu481300_mir); miR29c-3p (mmu479229_mir), miR29a-5p (mmu481032_mir), miR29b-5p (mmu481675_mir), miR29c-5p (mmu481034_mir).

### miRNA mimic and inhibition

miRNA overexpression or inhibition was carried out by transfection of 50 µM miRVANA miRNA mimic (4464066, Thermo Fisher) or 100 µM miRCURY LNA-inhibitor (YI04105459, Exiqon) for miR29c-5p or miR29c-3p. Transfections were carried out using Lipofectamine 2000 (11668030, Thermo Fisher) in Opti-MEM^™^ reduced serum media (31985070, Gibco) for DCs, and using Nucleofector electroporation kit (VPI-1001, Lonza) for exECs (Program M-003, Nucleofector 2b, Lonza).

### Luciferase assays

HEK-293 cells were co-transfected with 100 µM miR29c-5p miRVANA miRNA mimic (4464066, ThermoFisher) using Lipofectamine and one of the following Luciferase vectors, using RNAifectin (G073, Abm); Blank-Luc [pLenti-Ubc-UTR-Dual-Luc-Blank vector (C047, Abm)]; BMP4-3’UTR [pLenti-Ubc-3’UTR-Dual-Luciferase (MT-m02780-Custom)]; BMP4-5’UTR [pLenti-Ubc-Bmp4 (NM_001316360.1)-5’UTR-Dual Luciferase (C452, Abm)]; IL12B-3’UTR [pLenti-Ubc-IL12B-3’UTR-Dual-Luciferase (MT-10012-Custom)]; IL12B-5’UTR [pLenti-Ubc-IL12B-5’UTR-Dual-luciferase]; IL23A-3’UTR [pLenti-Ubc-Dual-Luciferase (MT-M10060-Custom)]; IL23A-5’UTR [pLenti-Ubc-IL23A-5’UTR-Dual-Luciferase (C452, Abm). Luciferase activity was measured after hours using the Dual-Glo® Luciferase Assay System (E2920, Promega) on a Veritas microplate Luminometer (Turner BioSystems, USA).

### Rho GTPase activation assays

Activated Rac1 was measured using the absorbance-based G-LISA Rac1 Activation Assay Biochem Kit (BK128, Cytoskeleton, USA). Briefly, exECs were co-cultured with thymocytes harvested from untreated mice, as described above. 24 hours after co-culture the exECs were harvested rapidly on ice, aliquoted and snap frozen using liquid nitrogen. Lysate volumes were subsequently adjusted for equal protein levels following BCA assay (23227, Pierce BCA protein assay kit, Thermo Fisher, USA), and GTP-bound Rac1 levels were assessed according to the manufacturers protocol. Plates were read at 490 nm on a Spark 10M plate reader (Tecan, Switzerland).

### Statistics

Statistical analysis between two groups was performed with unpaired two-tailed t test. Statistical comparison between 3 or more groups in Figs. 3f, 3h, 3k, and 4a was performed using a one-way ANOVA with Dunnett’s or Tukey’s multiple comparison test. Studies described in Figs 2c-f; Fig. 3a, 3b, 3i, 3j, and 3l used paired analyses. All statistics were calculated using Graphpad Prism and display graphs were generated in Graphpad Prism or R.

## Acknowledgements

We gratefully acknowledge the assistance of the Flow Cytometry and Comparative Medicine Core Facilities; and the support of the Immunotherapy Integrated Research Center at the Fred Hutchinson Cancer Research Center. We thank Dr. Brandon Hadland at the Fred Hutchinson Cancer Research Institute for generously providing the pCCL-PGK-myrAkt vector; and Dr. Marcel van den Brink (Memorial Sloan Kettering Cancer Center) for mentorship and discussions. This research was supported by National Institutes of Health award numbers R00-CA176376 (J.A.D.), R01-HL145276 (J.A.D.), Project 2 of P01-AG052359 (J.A.D.), and the NCI Cancer Center Support Grant P30-CA015704. Support was also received from a Scholar Award from the American Society of Hematology (J.A.D.), a Scholar Award from the Leukemia and Lymphoma Society (J.A.D.); the Mechtild Harf (John Hansen) Award from the DKMS Foundation for Giving Life (J.A.D.); the Cuyamaca Foundation (J.A.D.), and the Bezos Family Foundation (J.A.D.). S.K. was supported by a New Investigator Award from the American Society for Transplantation and Cellular Therapy and Pilot Funding from the Cooperative Center for Excellence in Hematology (Fred Hutchinson Cancer Research Center) award number U54 DK106829.

---

**Supplementary Figure 1.**
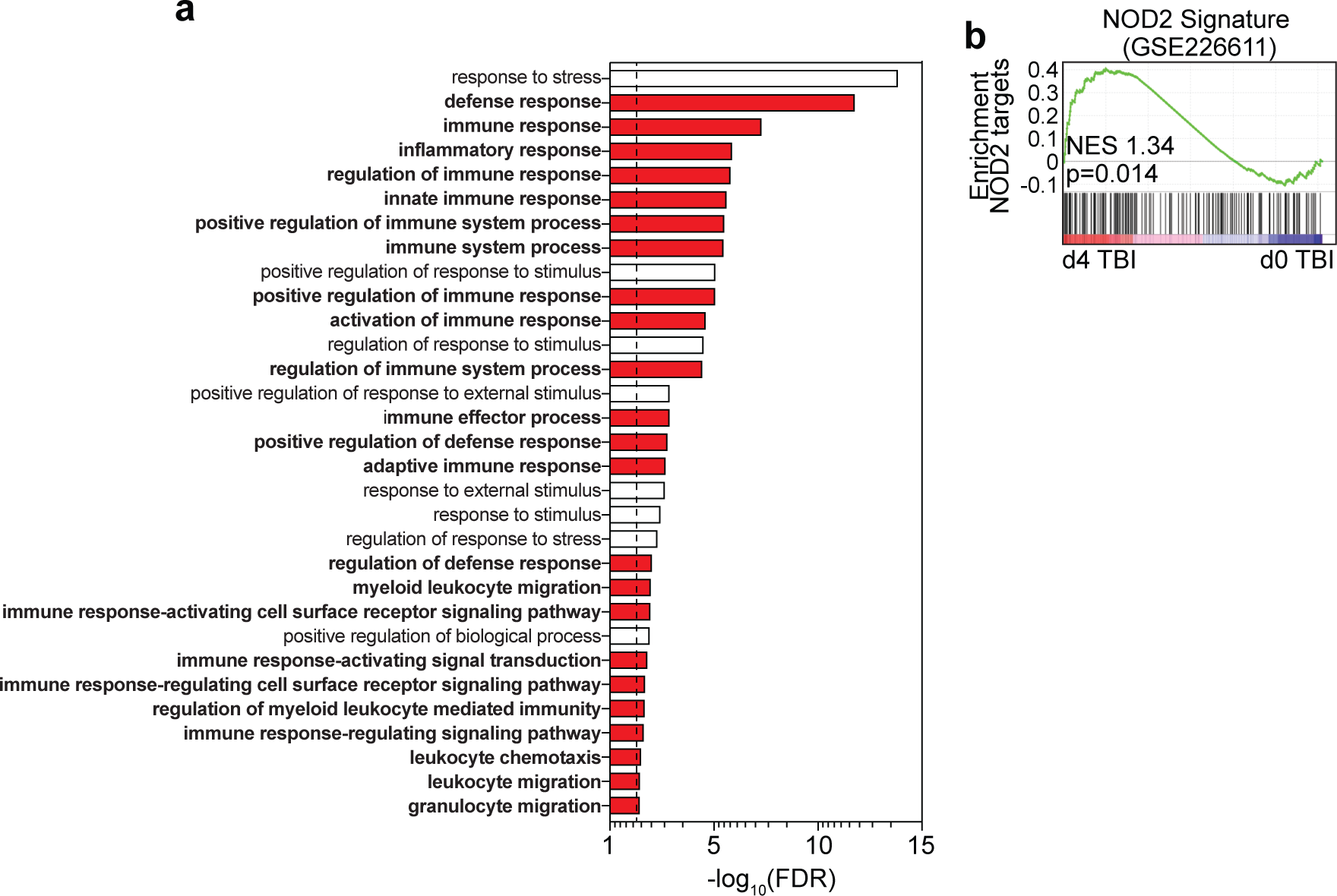
Thymuses were pooled from 6-week-old C57BL/6 mice and microarray analysis was performed on FACS purified ECs isolated from either untreated mice (d0) or 4 days after TBI (550 cGy, n=3/timepoint with each n pooled from 5 mice). **a**, Gene ontology (GO) pathway analysis was performed on upregulated genes at day 4 after SL-TBI using DAVID and all pathways with a FDR <0.05 are displayed. Red bars represent pathways involved with immune function. **b**, GSEA analysis was performed comparing gene expression changes at day 4 after SL-compared with a NOD2 gene signature (GSE226611). Heatmap showing gene expression changes at day 4 that match with the NOD2 gene signature.

**Supplementary Figure 2.**
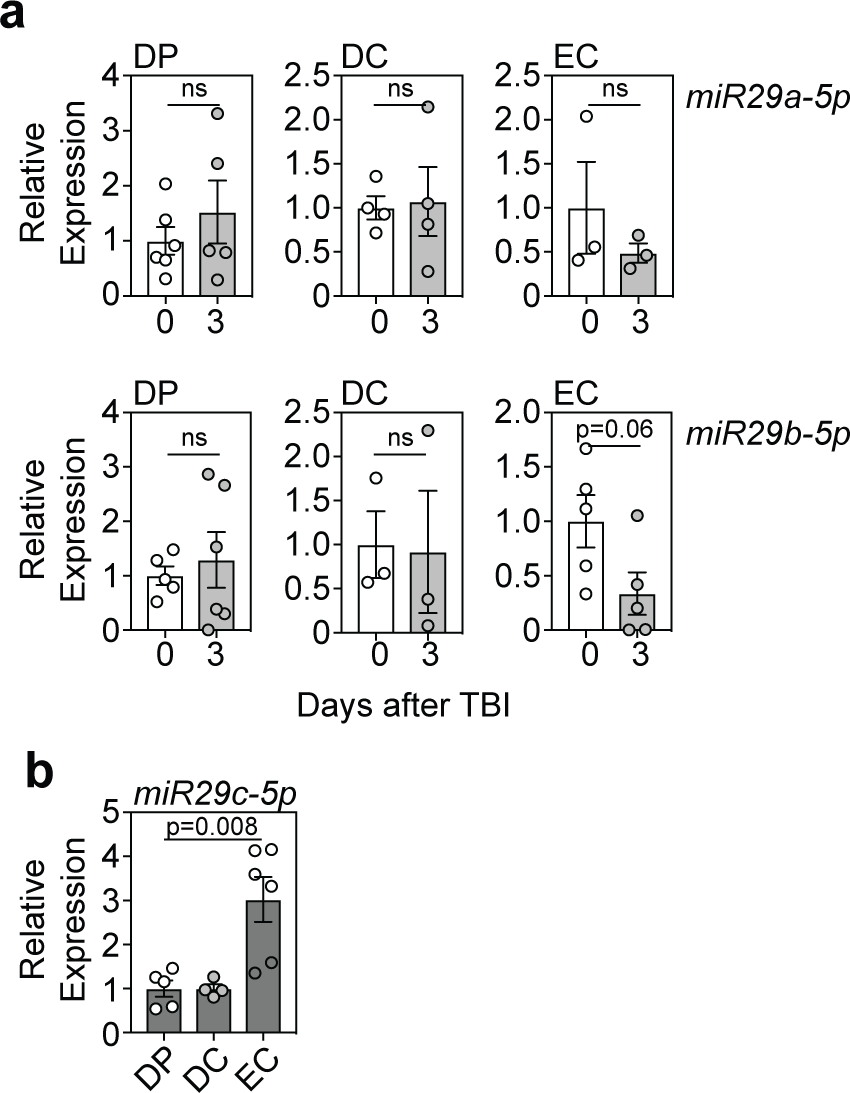
**a**, DPs, ECs and DCs were FACS purified from WT thymuses at day 0 or 3 after SL-TBI and expression of miR29a-5p and miR29b-5p was analyzed by qPCR (DP, n=5-6; DC, n=4; EC, n=3). **b**, DPs, ECs and DCs were FACS purified from WT thymuses at day 0 and expression of miR29c was analyzed by qPCR (DP, n=5; DC, n=4; EC, n=6). Graphs represent mean ± SEM, each dot represents a biologically independent observation.

**Supplementary Figure 3.**
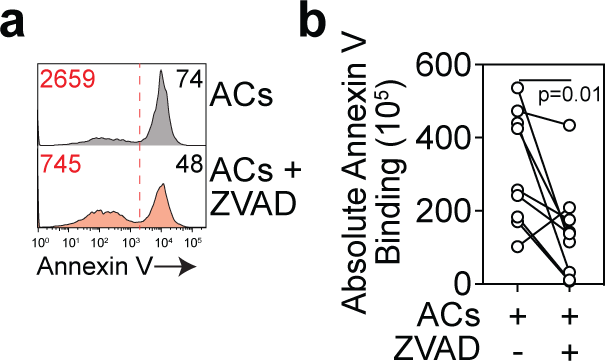
Thymocytes were isolated from untreated C57BL/6 mice and incubated for 4 hours with dexamethasone (100 nM) or zVAD-FMK (20 µM). After 4 hours, apoptotic thymocytes (ACs) were washed and co-cultured with exECs for 24 hours. **a**, Annexin V staining on CD4+CD8+ DP thymocytes (displayed are concatenated plots from 9 individual cultures, representative of three independent experiments). **b**, Absolute Annexin V binding in thymocytes used for co-culture experiments, measured at the end of the 4 h incubation period with dexamethasone (100 nM) or zVAD-FMK (20 µM), immediately prior to co-culture.

**Supplementary Figure 4.**
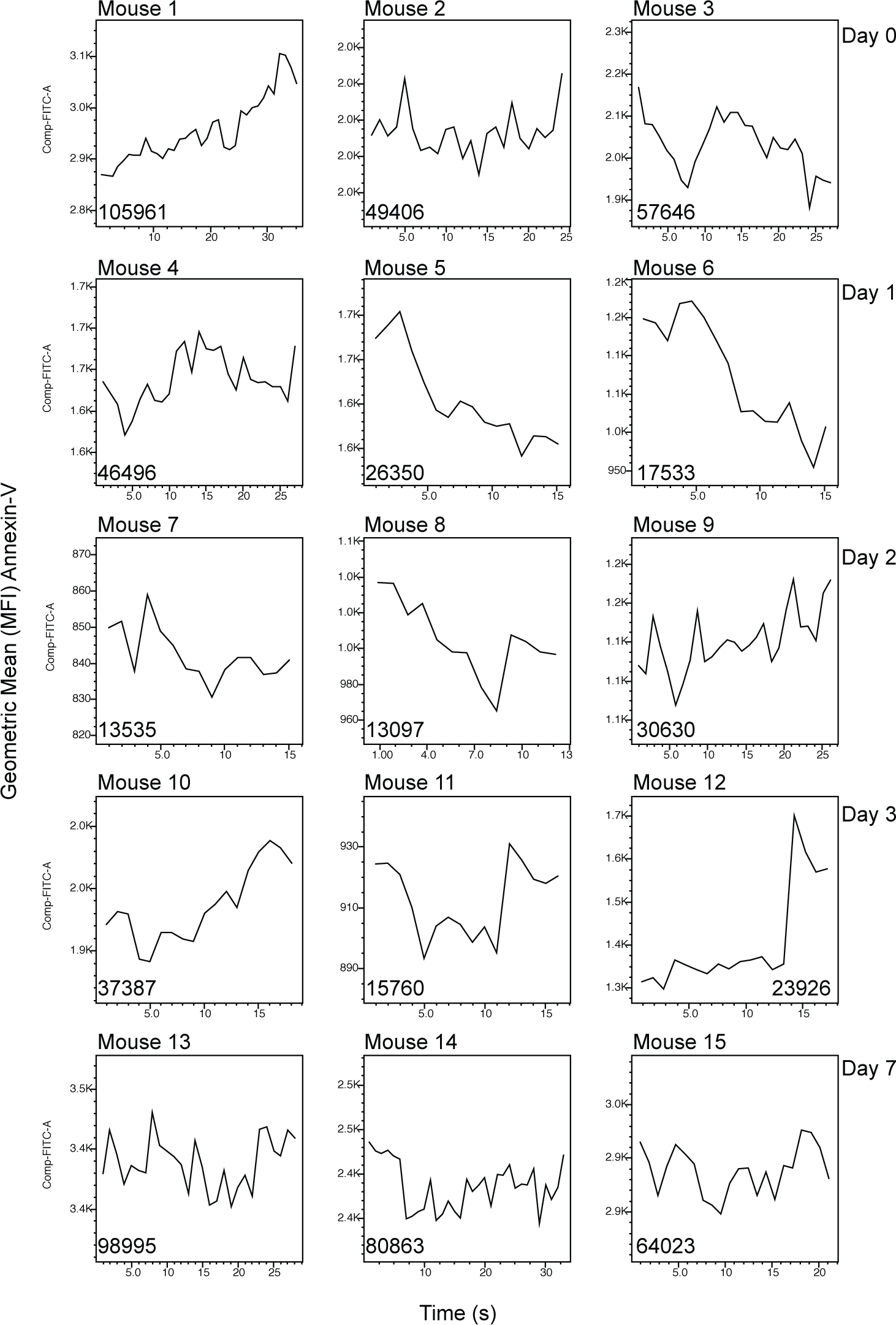
To calculate the absolute binding of Annexin V in the thymus, we first determined for each mouse at each time-point the calculation of the area under the curve (AUC) for the graph representing Geometric mean for Annexin V (gated only on singlets) across the time of the flow cytometry run. Absolute Annexin V binding in the thymus was then calculated by adjusting the total Annexin V binding for the flow run as a function of the total thymus cellularity for that individual mouse. Displayed are the AUC plots for each mouse from one of three experiments.

**Supplementary Figure 5.**
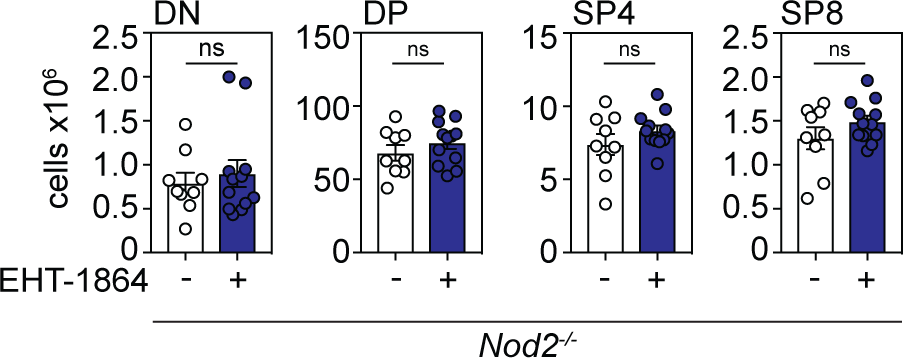
6-8 week Nod2-/- mice were treated with the Rac1 inhibitor EHT-1864 (40 mg/kg i.p. injection) at days 3, 5 and 7 following a sublethal dose of TBI (550 cGy). Total number of CD4+CD8+ DP, CD4+CD3+ SP4, or CD3+CD8+ SP8 thymocytes (n=9-12 across three independent experiments). Graphs represent mean ± SEM, each dot represents a biologically independent observation.

## References

1. Miller, J. The function of the thymus and its impact on modern medicine. Science (369(2020).

2. Abramson, J. & Anderson, G. Thymic Epithelial Cells. Annu Rev Immunol 35, 85–118 (2017).

3. Gruver, A.L. & Sempowski, G.D. Cytokines, leptin, and stress-induced thymic atrophy. Journal of Leukocyte Biology 84, 915–923 (2008).

4. van den Broek, T., et al. Neonatal thymectomy reveals differentiation and plasticity within human naive T cells. The Journal of clinical investigation 126, 1126–1136 (2016).

5. Kinsella, S. & Dudakov, J.A. When the Damage Is Done: Injury and Repair in Thymus Function. Frontiers in Immunology 11, 1745 (2020).

6. Chidgey, A., Dudakov, J., Seach, N. & Boyd, R. Impact of niche aging on thymic regeneration and immune reconstitution. Seminars in immunology 19, 331–340 (2007).

7. Merani, S., Pawelec, G., Kuchel, G.A. & McElhaney, J.E. Impact of Aging and Cytomegalovirus on Immunological Response to Influenza Vaccination and Infection. Frontiers in immunology 8, 784 (2017).

8. Bosch, M., Khan, F.M. & Storek, J. Immune reconstitution after hematopoietic cell transplantation. Curr Opin Hematol 19, 324–335 (2012).

9. van den Brink, M., Uhrberg, M., Jahn, L., DiPersio, J.F. & Pulsipher, M.A. Selected biological issues affecting relapse after stem cell transplantation: role of T-cell impairment, NK cells and intrinsic tumor resistance. Bone Marrow Transplant (2018).

10. Wertheimer, T., et al. Production of BMP4 by endothelial cells is crucial for endogenous thymic regeneration. Science Immunology (3(2018).

11. Dudakov, J.A., et al. Loss of thymic innate lymphoid cells leads to impaired thymopoiesis in experimental graft-versus-host disease. Blood 130, 933–942 (2017).

12. Dudakov, J.A., et al. Interleukin-22 Drives Endogenous Thymic Regeneration in Mice. Science 336, 91–95 (2012).

13. Buonocore, S., et al. Innate lymphoid cells drive interleukin-23-dependent innate intestinal pathology. Nature 464, 1371–1375 (2010).

14. Cella, M., et al. A human natural killer cell subset provides an innate source of IL-22 for mucosal immunity. Nature 457, 722–725 (2009).

15. Zheng, Y., et al. Interleukin-22, a TH17 cytokine, mediates IL-23-induced dermal inflammation and acanthosis. Nature 445, 648–651 (2007).

16. Caruso, R., Warner, N., Inohara, N. & Nunez, G. NOD1 and NOD2: signaling, host defense, and inflammatory disease. Immunity 41, 898–908 (2014).

17. Chen, G., Shaw, M.H., Kim, Y.G. & Nunez, G. NOD-like receptors: role in innate immunity and inflammatory disease. Annu Rev Pathol 4, 365–398 (2009).

18. Brain, O., et al. The Intracellular Sensor NOD2 Induces MicroRNA-29 Expression in Human Dendritic Cells to Limit IL-23 Release. Immunity 39, 521–536 (2013).

19. Kinnebrew, M.A., et al. Interleukin 23 Production by Intestinal CD103+CD11b+ Dendritic Cells in Response to Bacterial Flagellin Enhances Mucosal Innate Immune Defense. Immunity 36, 276–287 (2012).

20. Billmann-Born, S., et al. Genome-Wide Expression Profiling Identifies an Impairment of Negative Feedback Signals in the Crohn’s Disease-Associated NOD2 Variant L1007fsinsC. The Journal of Immunology 186, 4027–4038 (2011).

21. Iwanaga, Y., et al. Cloning, sequencing and expression analysis of the mouse NOD2/CARD15 gene. Inflammation research : official journal of the European Histamine Research Society … [et al.] 52, 272–276 (2003).

22. Martinic, M.M., et al. The Bacterial Peptidoglycan-Sensing Molecules NOD1 and NOD2 Promote CD8<sup>+</sup> Thymocyte Selection. The Journal of Immunology 198, 2649–2660 (2017).

23. Krol, J., Loedige, I. & Filipowicz, W. The widespread regulation of microRNA biogenesis, function and decay. Nat Rev Genet 11, 597–610 (2010).

24. Makeyev, E.V. & Maniatis, T. Multilevel regulation of gene expression by microRNAs. Science 319, 1789–1790 (2008).

25. Bartel, D.P. MicroRNAs: Genomics, Biogenesis, Mechanism, and Function. Cell 116, 281–297 (2004).

26. Ucar, O., Tykocinski, L.O., Dooley, J., Liston, A. & Kyewski, B. An evolutionarily conserved mutual interdependence between Aire and microRNAs in promiscuous gene expression. Eur J Immunol 43, 1769–1778 (2013).

27. Papadopoulou, A.S., et al. The thymic epithelial microRNA network elevates the threshold for infection-associated thymic involution via miR-29a mediated suppression of the IFN-[alpha] receptor. Nat Immunol 13, 181–187 (2012).

28. Ye, Y., et al. MicroRNA expression in the aging mouse thymus. Gene 547, 218–225 (2014).

29. Seandel, M., et al. Generation of a functional and durable vascular niche by the adenoviral E4ORF1 gene. Proceedings of the National Academy of Sciences of the United States of America 105, 19288–19293 (2008).

30. Huntzinger, E. & Izaurralde, E. Gene silencing by microRNAs: contributions of translational repression and mRNA decay. Nat Rev Genet 12, 99–110 (2011).

31. Mak, T.W., Penninger, J.M. & Ohashi, P.S. Knockout mice: a paradigm shift in modern immunology. Nat Rev Immunol 1, 11–19 (2001).

32. Purton, J.F., et al. Expression of the glucocorticoid receptor from the 1A promoter correlates with T lymphocyte sensitivity to glucocorticoid-induced cell death. J Immunol 173, 3816–3824 (2004).

33. Jameson, S.C., Hogquist, K.A. & Bevan, M.J. Positive selection of thymocytes. Annu Rev Immunol 13, 93–126 (1995).

34. Hernandez, J.B., Newton, R.H. & Walsh, C.M. Life and death in the thymus--cell death signaling during T cell development. Curr Opin Cell Biol 22, 865–871 (2010).

35. Municio, C., et al. Apoptotic cells enhance IL-10 and reduce IL-23 production in human dendritic cells treated with zymosan. Molecular Immunology 49, 97–106 (2011).

36. Cifone, M.G., et al. Dexamethasone-Induced Thymocyte Apoptosis: Apoptotic Signal Involves the Sequential Activation of Phosphoinositide-Specific Phospholipase C, Acidic Sphingomyelinase, and Caspases. Blood 93, 2282–2296 (1999).

37. Mower, D.A., Jr., et al. Decreased membrane phospholipid packing and decreased cell size precede DNA cleavage in mature mouse B cell apoptosis. J Immunol 152, 4832–4842 (1994).

38. Schlegel, R.A., Stevens, M., Lumley-Sapanski, K. & Williamson, P. Altered lipid packing identifies apoptotic thymocytes. Immunology letters 36, 283–288 (1993).

39. Wallet, M.A., et al. MerTK regulates thymic selection of autoreactive T cells. Proceedings of the National Academy of Sciences of the United States of America 106, 4810–4815 (2009).

40. Rothlin, C.V., Carrera-Silva, E.A., Bosurgi, L. & Ghosh, S. TAM receptor signaling in immune homeostasis. Annu Rev Immunol 33, 355–391 (2015).

41. Lemke, G. & Rothlin, C.V. Immunobiology of the TAM receptors. Nat Rev Immunol 8, 327–336 (2008).

42. Yokoyama, Y., et al. Immuno-Oncological Efficacy of RXDX-106, a Novel Small Molecule Inhibitor of the TAM (TYRO3, AXL, MER) Family of Kinases. Cancer Research, canres.2022.2018 (2019).

43. Keestra-Gounder, A.M. & Tsolis, R.M. NOD1 and NOD2: Beyond Peptidoglycan Sensing. Trends Immunol 38, 758–767 (2017).

44. Keestra, A.M., et al. Manipulation of small Rho GTPases is a pathogen-induced process detected by NOD1. Nature 496, 233–237 (2013).

45. Herve, J.C. & Bourmeyster, N. Rho GTPases at the crossroad of signaling networks in mammals. Small GTPases 6, 43–48 (2015).

46. Cleverley, S., Henning, S. & Cantrell, D. Inhibition of Rho at different stages of thymocyte development gives different perspectives on Rho function. Curr Biol 9, 657–660 (1999).

47. Gomez, M., Kioussis, D. & Cantrell, D.A. The GTPase Rac-1 controls cell fate in the thymus by diverting thymocytes from positive to negative selection. Immunity 15, 703–713 (2001).

48. Gomez, M., Tybulewicz, V. & Cantrell, D.A. Control of pre-T cell proliferation and differentiation by the GTPase Rac-I. Nat Immunol 1, 348–352 (2000).

49. Hodge, R.G. & Ridley, A.J. Regulating Rho GTPases and their regulators. Nat Rev Mol Cell Biol 17, 496–510 (2016).

50. Lin, Y. & Zheng, Y. Approaches of targeting Rho GTPases in cancer drug discovery. Expert Opin Drug Discov 10, 991–1010 (2015).

51. Mazieres, J., Pradines, A. & Favre, G. Perspectives on farnesyl transferase inhibitors in cancer therapy. Cancer Lett 206, 159–167 (2004).

52. Shang, X., et al. Rational design of small molecule inhibitors targeting RhoA subfamily Rho GTPases. Chem Biol 19, 699–710 (2012).

53. Shutes, A., et al. Specificity and mechanism of action of EHT 1864, a novel small molecule inhibitor of Rac family small GTPases. The Journal of biological chemistry 282, 35666–35678 (2007).

54. Dong, M., et al. Rho-kinase inhibition: a novel therapeutic target for the treatment of cardiovascular diseases. Drug Discov Today 15, 622–629 (2010).

55. Sadok, A., et al. Rho kinase inhibitors block melanoma cell migration and inhibit metastasis. Cancer Res 75, 2272–2284 (2015).

56. Goldberg, G.L., et al. Sex Steroid Ablation Enhances Immune Reconstitution Following Cytotoxic Antineoplastic Therapy in Young Mice. J Immunol 184, 6014–6024 (2010).

57. Dudakov, J.A., Perales, M.A. & van den Brink, M.R.M. Immune Reconstitution Following Hematopoietic Cell Transplantation. in Thomas’ Hematopoietic Cell Transplantation, Vol. 1 (eds. Forman, S., Negrin, R.S., Antin, J.H. & Appelbaum, F.A.) 160–165 (John Wiley & Sons, Ltd., West Sussex, UK, 2016).

58. Mackall, C.L., et al. Age, thymopoiesis, and CD4+ T-lymphocyte regeneration after intensive chemotherapy. N Engl J Med 332, 143–149 (1995).

59. Parkman, R. & Weinberg, K.I. Immunological reconstitution following bone marrow transplantation. Immunol Rev 157, 73–78 (1997).

60. Weinberg, K., et al. The effect of thymic function on immunocompetence following bone marrow transplantation. Biol Blood Marrow Transplant 1, 18–23 (1995).

61. Komanduri, K.V., et al. Delayed immune reconstitution after cord blood transplantation is characterized by impaired thymopoiesis and late memory T-cell skewing. Blood 110, 4543–4551 (2007).

62. Legrand, N., Dontje, W., van Lent, A.U., Spits, H. & Blom, B. Human thymus regeneration and T cell reconstitution. Semin Immunol 19, 280–288 (2007).

63. Mackall, C.L. T-cell immunodeficiency following cytotoxic antineoplastic therapy: a review. Stem Cells 18, 10–18 (2000).

64. Pizzo, P.A., Rubin, M., Freifeld, A. & Walsh, T.J. The child with cancer and infection. II. Nonbacterial infections. The Journal of pediatrics 119, 845–857 (1991).

65. Williams, K.M., Hakim, F.T. & Gress, R.E. T cell immune reconstitution following lymphodepletion. Semin Immunol 19, 318–330 (2007).

66. Clave, E., et al. Thymic function recovery after unrelated donor cord blood or T-cell depleted HLA-haploidentical stem cell transplantation correlates with leukemia relapse. Front Immunol 4, 54 (2013).

67. Jaffe, H.L. The Influence of the Suprarenal Gland on the Thymus : I. Regeneration of the Thymus Following Double Suprarenalectomy in the Rat. J Exp Med 40, 325–342 (1924).

68. Miller, J.F. Immunological function of the thymus. Lancet 2, 748–749 (1961).

69. Dudakov, J.A., Khong, D.M.P., Boyd, R.L. & Chidgey, A.P. Feeding the fire: the role of defective bone marrow function in exacerbating thymic involution. Trends Immunol 31, 191–198 (2010).

70. Martinic, M.M., et al. The Bacterial Peptidoglycan-Sensing Molecules NOD1 and NOD2 Promote CD8(+) Thymocyte Selection. J Immunol 198, 2649–2660 (2017).

71. Watanabe, T., Kitani, A., Murray, P.J. & Strober, W. NOD2 is a negative regulator of Toll-like receptor 2-mediated T helper type 1 responses. Nat Immunol 5, 800–808 (2004).

72. Matsuo, M., et al. MiR-29c is downregulated in gastric carcinomas and regulates cell proliferation by targeting RCC2. Mol Cancer 12, 15 (2013).

73. Watts, A.E., et al. MicroRNA29a Treatment Improves Early Tendon Injury. Mol Ther 25, 2415–2426 (2017).

74. Chen, L., et al. Modulation of miR29a improves impaired post-ischemic angiogenesis in hyperglycemia. Experimental biology and medicine (Maywood, N.J 242, 1432–1443 (2017).

75. Mersey, B.D., Jin, P. & Danner, D.J. Human microRNA (miR29b) expression controls the amount of branched chain alpha-ketoacid dehydrogenase complex in a cell. Human molecular genetics 14, 3371–3377 (2005).

76. Batliner, J., et al. Transcriptional regulation of MIR29B by PU.1 (SPI1) and MYC during neutrophil differentiation of acute promyelocytic leukaemia cells. Br J Haematol 157, 270–274 (2012).

77. Wang, L.H., et al. Downregulation of miR29b targets DNMT3b to suppress cellular apoptosis and enhance proliferation in pancreatic cancer. Molecular medicine reports 17, 2113–2120 (2018).

78. Chen, B., et al. Inhibition of miR-29c promotes proliferation, and inhibits apoptosis and differentiation in P19 embryonic carcinoma cells. Molecular medicine reports 13, 2527–2535 (2016).

79. Municio, C., et al. Apoptotic cells enhance IL-10 and reduce IL-23 production in human dendritic cells treated with zymosan. Mol Immunol 49, 97–106 (2011).

80. Strober, W., Murray, P.J., Kitani, A. & Watanabe, T. Signalling pathways and molecular interactions of NOD1 and NOD2. Nat Rev Immunol 6, 9–20 (2006).

81. Nunes-Alves, C., Nobrega, C., Behar, S.M. & Correia-Neves, M. Tolerance has its limits: how the thymus copes with infection. Trends in immunology 34, 502–510 (2013).

82. Eitel, J., et al. Beta-PIX and Rac1 GTPase mediate trafficking and negative regulation of NOD2. J Immunol 181, 2664–2671 (2008).

83. Keestra, A.M. & Baumler, A.J. Detection of enteric pathogens by the nodosome. Trends Immunol 35, 123–130 (2014).

84. Saxena, M. & Yeretssian, G. NOD-Like Receptors: Master Regulators of Inflammation and Cancer. Front Immunol 5, 327 (2014).

85. Legrand-Poels, S., et al. Modulation of Nod2-dependent NF-kappaB signaling by the actin cytoskeleton. J Cell Sci 120, 1299–1310 (2007).

86. Wennerberg, K. & Der, C.J. Rho-family GTPases: it’s not only Rac and Rho (and I like it). J Cell Sci 117, 1301–1312 (2004).

87. Van Aelst, L. & D’Souza-Schorey, C. Rho GTPases and signaling networks. Genes & development 11, 2295–2322 (1997).

88. Krieser, R.J. & Eastman, A. Cleavage and nuclear translocation of the caspase 3 substrate Rho GDP-dissociation inhibitor, D4-GDI, during apoptosis. Cell Death Differ 6, 412–419 (1999).

89. Na, S., et al. D4-GDI, a substrate of CPP32, is proteolyzed during Fas-induced apoptosis. The Journal of biological chemistry 271, 11209–11213 (1996).

90. Coleman, M.L. & Olson, M.F. Rho GTPase signalling pathways in the morphological changes associated with apoptosis. Cell Death Differ 9, 493–504 (2002).

91. Koyanagi, M., et al. Inhibition of the Rho/ROCK pathway reduces apoptosis during transplantation of embryonic stem cell-derived neural precursors. Journal of neuroscience research 86, 270–280 (2008).

92. Lacal, J.C. Regulation of proliferation and apoptosis by Ras and Rho GTPases through specific phospholipid-dependent signaling. FEBS Letters 410, 73–77 (1997).

93. Sanno, H., et al. Control of postnatal apoptosis in the neocortex by RhoA-subfamily GTPases determines neuronal density. J Neurosci 30, 4221–4231 (2010).

94. Dumont, C., et al. Rac GTPases play critical roles in early T-cell development. Blood 113, 3990–3998 (2009).

95. Cicin-Sain, L., et al. Loss of naive T cells and repertoire constriction predict poor response to vaccination in old primates. Journal of immunology (Baltimore, Md. : 1950) 184, 6739–6745 (2010).

96. Zhang, N., Hartig, H., Dzhagalov, I., Draper, D. & He, Y.W. The role of apoptosis in the development and function of T lymphocytes. Cell Res 15, 749–769 (2005).

97. Guo, F., Cancelas, J.A., Hildeman, D., Williams, D.A. & Zheng, Y. Rac GTPase isoforms Rac1 and Rac2 play a redundant and crucial role in T-cell development. Blood 112, 1767–1775 (2008).

98. Velardi, E., et al. Sex steroid blockade enhances thymopoiesis by modulating Notch signaling. The Journal of Experimental Medicine 211, 2341–2349 (2014).

99. Huang da, W., Sherman, B.T. & Lempicki, R.A. Systematic and integrative analysis of large gene lists using DAVID bioinformatics resources. Nat Protoc 4, 44–57 (2009).

100. Huang da, W., Sherman, B.T. & Lempicki, R.A. Bioinformatics enrichment tools: paths toward the comprehensive functional analysis of large gene lists. Nucleic Acids Res 37, 1–13 (2009).

